# Non-invasive estrous cycle classification in mice using convolutional neural networks

**DOI:** 10.1101/2025.10.03.680273

**Authors:** Yassin Terki, Mohiuddin Saifullah, Gema Puspa Sari, Ayako Isotani, Yuichi Sakumura

## Abstract

Accurately identifying the estrous cycle phases in laboratory mice is essential for neurological research, reproductive studies, and breeding programs. Here, we introduce a non-invasive approach using Convolutional Neural Networks (CNNs) to classify estrous stages from external genital images, with cytological smears serving as the reference standard for labeling. Images were curated, preprocessed, and augmented to ensure consistency and generalization. Seven pre-trained CNNs and a lightweight custom architecture, Repro Cycle Net (RCN), were evaluated. All models achieved accuracies above 75%, with RCN exceeding 83% and showing the lowest test loss, highlighting its efficiency despite a simple four-layer design. Saliency map analyses revealed that classification relied on perivaginal features, while irrelevant regions such as the fur area were largely ignored. Importantly, binary classification of estrus versus non-estrus directly informs mating feasibility, underscoring the immediate utility of this method for reproductive studies and colony management. This work demonstrates that combining deep learning with external genital observation enables efficient and reproducible estrous monitoring, supporting both experimental reliability and animal welfare.

## Introduction

In mammalian developmental studies, understanding the estrous cycle in female mice is essential, as hormonal fluctuations during this cycle profoundly influence tissue development, physiology, and behavior. Properly accounting for these fluctuations is critical for improving the accuracy and reproducibility of experimental outcomes [1]. Despite the predominance of male mice in disease model analysis, incorporating female mice is necessary to ensure a comprehensive understanding of behavioral analysis, including the issue of sex differences in experimental studies [1,2]. The estrous cycle, similar to the human menstrual cycle, involves ovarian and uterine changes that affect a wide range of biological processes, from genetic engineering in animal breeding to behavioral and physiological studies [3,10]. However, accurately determining the estrous cycle in female mice remains a significant challenge. This limitation reduces experimental reliability and hinders the study of biological processes. Resolving this issue would not only enhance reproducibility but also contribute to a more comprehensive understanding of developmental biology.

Several methods exist for determining the estrous cycle in female mice. One approach involves hormone measurement, where the stage of the cycle is identified by quantifying the concentrations of estrogen and progesterone in serum and urine [3]. While this method is highly accurate, it requires frequent sampling, which may burden the experimental animals [3]. Another approach is behavioral observation, where increases in specific exploratory and social behaviors during the estrous period serve as indicators [4]. Although non-invasive, this method relies on subtle behavioral changes, requiring an experienced observer [4]. Additionally, cytological examination through the vaginal smear method is widely used, with the stage of the cycle identified by collecting and microscopically examining vaginal cells [5]. However, this method can cause some stress to the animals and requires technical skill [5]. Lastly, visual observation—a simple method that involves checking for swelling along with color changes in the external genitalia including the presence of moist and appearance of numerous longitudinal folds or striations—is highly subjective and has limited accuracy [6].

Despite these efforts, there is still no non-invasive and fully objective method that allows reliable and automated staging of the estrous cycle in laboratory mice. This research gap is critical because subjective or invasive approaches inevitably reduce reproducibility and animal welfare.

Recent advancements in machine learning, particularly deep learning, are expected to improve the accuracy and efficiency of estrous cycle determination. Image recognition technology, in particular, holds promise for reducing subjective judgments and enabling consistent, highly accurate analyses. Indeed, deep learning techniques using vaginal cell images have been developed and have demonstrated high accuracy in classifying estrous cycles [7]. However, the requirement to collect vaginal cells and the associated burden on the animals remains a concern. Therefore, a method that enables estrous cycle determination directly from external genital images would provide a practical, non-invasive, and animal welfare–conscious alternative [8].

In this study, we employed convolutional neural networks (CNN), a deep learning method, to determine the estrous cycle based on images of the external genitalia of female mice. The labels for the images were determined using the vaginal smear method to identify each estrous stage, which served as a reference [5,9]. The dataset was preprocessed by adjusting the position of the vagina in the images and correcting for brightness. Using this dataset, we evaluated several pretraining models along with our own developed model and achieved an accuracy of over 75%. The overall workflow of this study, including preprocessing, feature extraction, classification, and model validation, is summarized in **Fig. 1**. In addition to testing widely used pretrained CNNs, we designed a lightweight custom architecture—Repro Cycle Net (RCN)—specifically optimized for estrous classification tasks. Our results demonstrate, for the first time, that external genital images alone can provide sufficient features for reliable estrous classification, without the need for invasive cytological sampling. These results demonstrate that combining machine learning with visual observation of external genitalia enables efficient and accurate determination of the estrous cycle in female mice. This approach may serve as a practical and reproducible standard for estrous monitoring in laboratory mice, supporting both experimental reliability and animal welfare. In particular, a binary classification of estrous versus non-estrous directly informs mating feasibility, which is highly relevant for reproductive studies and colony management.

**Figure 1.**
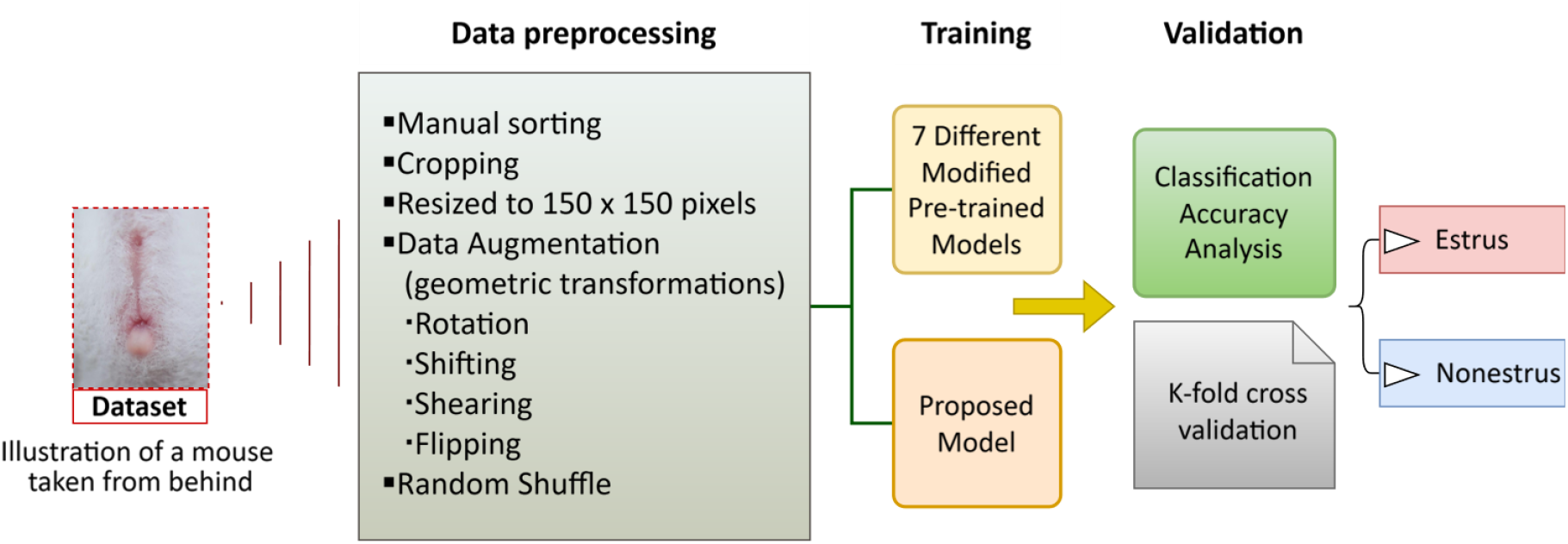
Workflow of the different associative steps including preprocessing, feature extraction, classification, model validation.

## Materials & Methods

### Mice and animal guidelines

Animal handling, breeding, and all experimental procedures were conducted under specific-pathogen-free (SPF) conditions at the Nara Institute of Science and Technology (NAIST), according to the guidelines of “Regulations and By-Laws of Animal Experimentation at the Nara Institute for Science and Technology”, and approved by the Animal experimental Committee (approval no. 1639 and no. 2103), and carried out in compliance with the ARRIVE guidelines [11]. Albino ICR mice used in this study were purchased from SLC, Japan, and maintained in the NAIST Animal Facility. The mice were maintained under a 12-hours light/ 12-hours dark cycle (lights on 08:00 am). The maximum caging density for female mice was five mice from the same litter and sex starting from weaning. Female ages in this study were 7-18 weeks. Male vasectomized mice were prepared by cutting their vas deferens and were used to induce pseudopregnancy in foster mothers for embryo transfer of genetically manipulated fertilized embryo. Food and water are available at libitum.

### Visual examination of the estrous stage

Visual examination of the estrous stage was performed based on criteria described by Champlin [20]. Female mice were placed on the wire grid and gently restrained by their front paws. By holding their tails, the rear end of the females was lifted slightly above the grid toward the observer. The vagina was observed under normal fluorescent room lighting, and the appearance was photographed with a digital camera (CANON Powershot G9X Mark II; resolution 5472 × 3648 pixels) at a distance of approximately 20–40 cm (**Fig. 2**). Visual examination and image acquisition were performed at 08:00–11:00 am and 02:00–04:00 pm. These time windows were chosen to minimize variability due to circadian changes and to ensure consistent lighting conditions.

**Figure 2.**
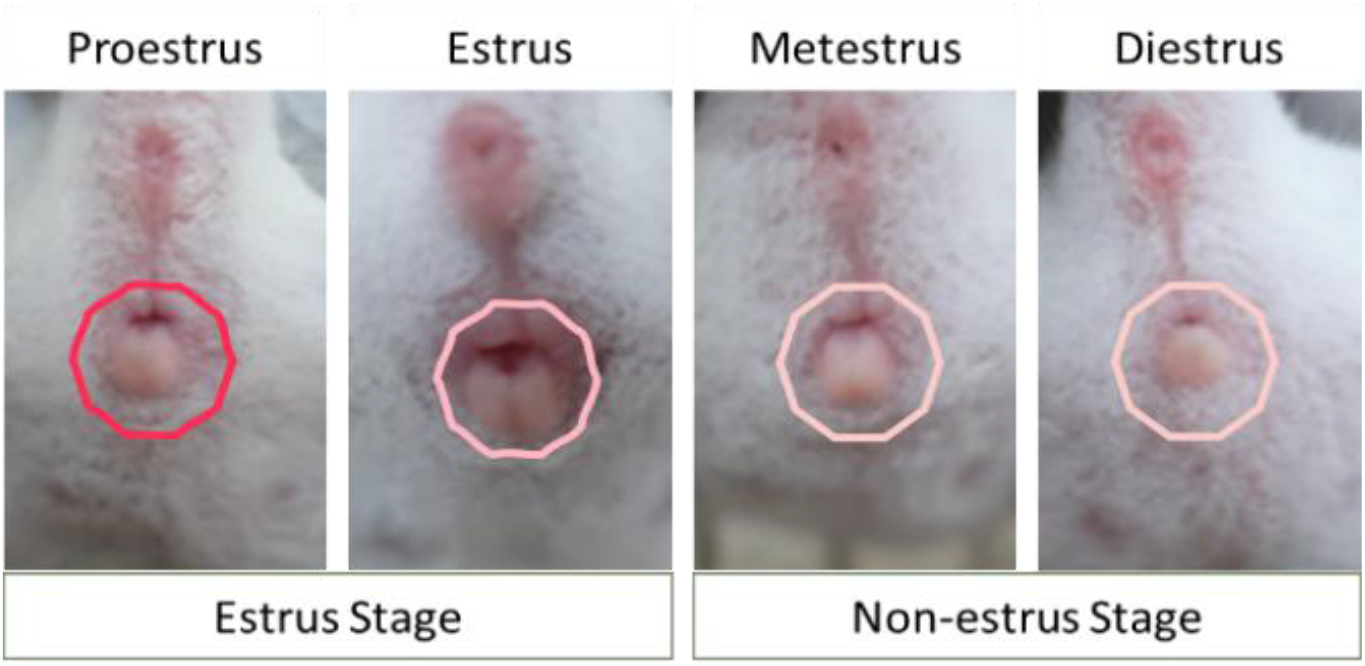
Visual assessment used to identify the estrous cycle in female mice. The four biological stages (proestrus, estrus, metestrus, and diestrus) were grouped and labeled as either estrous or non-estrous for binary classification.

Cytological examination of the estrous stage performed as described by Byers et al. (2012) [21] with modification on cell transfer process to the glass slide. Vaginal cells were collected by swab method using a small size cotton-tipped wetted with room temperature phosphate buffer saline (PBS, Sigma) and inserted into the vagina of the restrained mouse. The cotton tip then turned and rolled carefully against the wall of the vagina. The cotton tip was then dipped into 25 ul PBS (Sigma). The vaginal cells were then transferred into a glass slide (Matsunami micro slide glass S2112), and then dried inside a 56°C oven for 20 minutes, and then stained with crystal violet (Merck 109218) for 90 seconds. The slides were washed carefully with tap water to rinse the excess staining. After drying, microscopic observation was done under a light microscope with 20 x magnification to observe all cell types. The cell types in different stages of the estrous cycle were determined with the focus on the cell types that were characteristic of the different estrous stages [12]. Samples were collected from 08:00 am - 11:00 am.

### Data preprocessing

The raw data obtained for this study required preprocessing to ensure its suitability for training the deep learning models. The following steps outline the preprocessing pipeline executed:

1. To resolve image quality issues, such as out-of-focus or poorly illuminated images, we first sorted the raw dataset and excluded unsuitable samples, prioritizing three key parameters: vaginal swelling, color, and opening.
2. Due to these parameters, we grouped the original four estrous stages into only two classes (estrous and non-estrous; **Fig. 2**). This binary classification was chosen to simplify the analysis and to capture the most biologically relevant contrast, namely whether estrogen is exerting physiological effects or not, rather than overfitting to fine-grained stage distinctions. The distribution of images across the original four stages and their grouping into the two final classes is summarized in **Table 1**.
3. Subsequently, we manually split the dataset into training and testing subsets with an 90:10 ratio to evaluate the model’s performance effectively.
4. To confirm a consistent input dimension for the deep learning model, all images were resized to a fixed size of 150 x 150 pixels using the appropriate libraries across all implemented models.
5. To expand the diversity of the training dataset and enhance the model’s generalization capabilities, we implemented data augmentation while carefully preserving the original color fidelity. Our augmentation primarily involved geometric transformations, including rotation, shifting, shearing, and flipping. Minor color variations (e.g., a slight bluish hue) sometimes appeared due to interpolation, but these artifacts had no significant impact on dataset integrity. During the model training process, we utilized both raw and augmented data from each class to ensure robust learning.
6. Finally, the dataset was shuffled to prior training to introduce randomness and minimize bias

**Table 1.** Distribution of images across the four estrous stages and their grouping into two final classes (estrus and non-estrus) after preprocessing. ‘Merged’ indicates the combined image counts per final class.

| Stage | Data type | Raw images | Sorted images | Merged (per class) | Final class |
| --- | --- | --- | --- | --- | --- |
| Proestrus | RGB Image | 250 | 193 | 403 | **Estrus** |
| Estrus |  | 256 | 210 |  |  |
| Metestrus |  | 147 | 87 | 355 | **Non-estrus** |
| Diestrus |  | 356 | 268 |  |  |

These preprocessing steps secured the dataset that was suitably formatted and enriched with diverse samples, thereby enhancing the model’s ability to learn discriminative features and generalize well to unseen new data.

### Models and analysis

#### Existing models

To extract the feature, we applied seven pre-trained models along with our own proposed model. Transfer learning entails leveraging a pre-trained model, initially trained on a particular problem, for a similar task. This approach offers the benefit of reduced training time, as the model has already learned relevant features. In the context of image classification, many CNN models, previously validated in the ImageNet challenge, serve as pre-trained models. In this study, we have applied seven pre-trained CNN models: Densenet [13], EfficientNet [14], Inception [15], Mobilenet [16], NasNet [17], ResNet [18], & Xception [19]. For transfer learning, the convolutional layers of each model were used as fixed feature extractors, while the fully connected layers were re-trained using our dataset. Classification was performed with each of these adapted models, enabling a direct performance comparison with our proposed model. To mitigate overfitting, key hyperparameters such as learning rate, dropout, and dense layer size were tuned using random search techniques.

#### Proposed model

The proposed Repro Cycle Net (RCN) is a model specifically designed and adjusted to suit our specific purposes. The model was aimed to ensure precision and efficiency in computation and to compare its performance with the pretrained ones, thereby providing greater significance to this analysis. The architecture of the RCN model consists of four distinct convolutional layers, structured into one single convolutional layer followed by two consecutive convolutional layers and a final deeper layer (**Fig. 3**). Each layer is configured with specific filter sizes, with one layer set at 16 filters, two layers set at 32 filters and the final one at 64 filters. Each convolutional layer was followed by a ReLU activation and max-pooling operation, after which the extracted features were flattened and passed to a fully connected dense layer with dropout regularization. The classifier consisted 2 layer which include one layer of 256 neurons with a 0.5 dropout rate and a layer of a single unit for binary classification. This configuration enhances the model’s ability to process images, capturing intricate patterns with increased detail and accuracy while maintaining a lightweight design.

**Figure 3.**
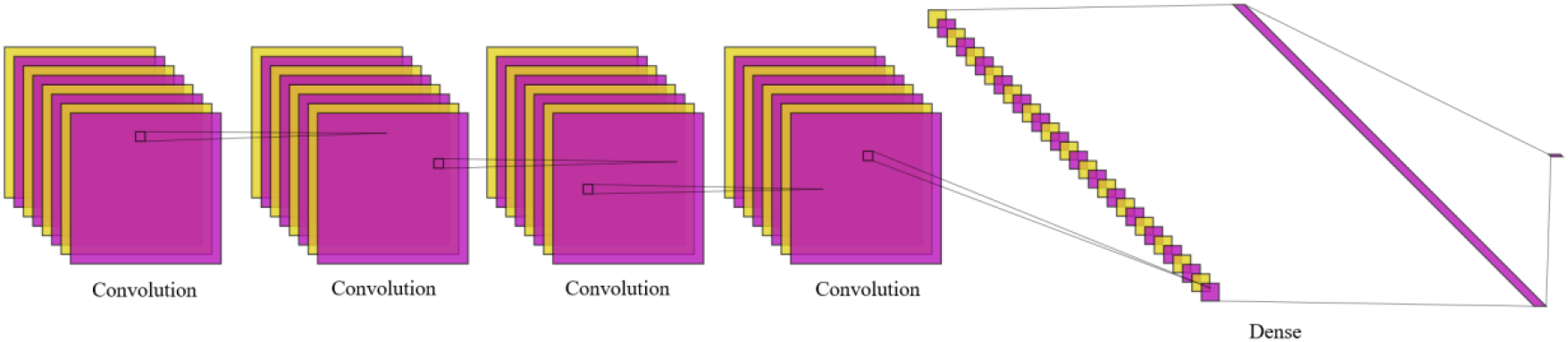
Architecture of the proposed Repro Cycle Net (RCN). The model consists of four convolutional layers (16, 32, 32, and 64 filters), each followed by ReLU activation and max pooling, a fully connected dense layer with dropout, and a final sigmoid output layer for binary classification.

Beyond architectural considerations, our model development process involved extensive parameter calibration. We fine-tuned various hyperparameters, including learning rates (explored in the range of 10^−5^ to 10^−3^), initializers, dropout rate (0.5), batch size (64), pruning strategies, and methods to prevent overfitting. Pruning was applied during training to remove redundant connections and improve computational efficiency. These adjustments were essential to align the model with the specific requirements of our research objectives and are documented in the provided codebase.

#### Performance evaluation, feature extraction, programming, and computation

In order to evaluate the classification accuracy of the model in estimating the estrous cycle, we introduced the indices of classification accuracy, test loss, and ROC-AUC. Classification accuracy was defined as the accuracy when the threshold of the output layer of the model was set to 0.5, calculated as the ratio of correctly classified samples (true positives + true negatives) to the total number of samples. The test loss was calculated using the binary cross-entropy loss function. The ROC curve was created by changing the discrimination threshold of the output layer of the model from 0 to 1, and the corresponding area under the curve (AUC) was used as a threshold-independent performance metric.

Furthermore, we used the saliency map method, specifically a vanilla gradient-based saliency approach, to illustrate the important pixels the model is paying attention to. This analysis enabled us to identify which areas of the images the model relied on for classification and whether these regions corresponded to biologically relevant objectives and preferred positions.

The development of both the existing models and the RCN was conducted using the Python programming language and TensorFlow library’s Keras interface. Computations were performed on a GPU computer (NVIDIA GeForce RTX 4070 Ti, 4 GB VRAM, CUDA 11.2, 128 GB system RAM).

## Results

We trained seven pre-trained models along with our independently developed RCN using learning images of the vaginal region of female mice. Subsequently, we evaluated binary classification accuracy (estrus vs. non-estrus) and test loss using test images (**Fig. 4A**). Higher classification accuracy indicates a more precise determination of estrous, while lower test loss suggests a greater difference in the probability used as the classification threshold.

**Figure 4.**
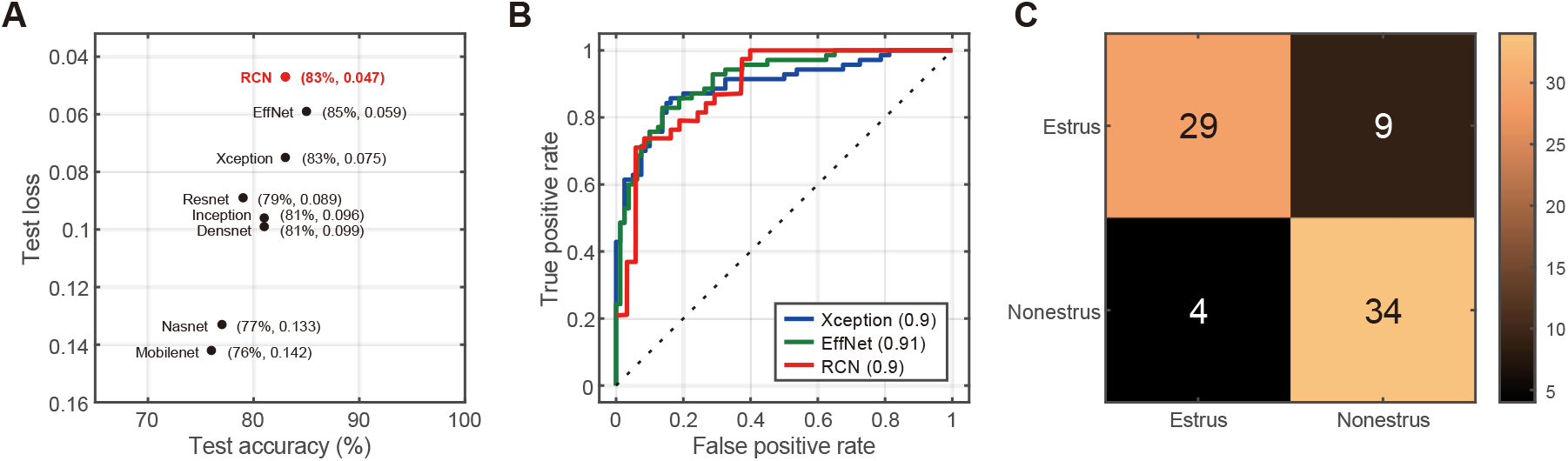
Performance comparison of CNN models for estrous classification. (**A**) Comparison of the seven pretrained models including the RCN. The horizontal axis represents classification accuracy, while the vertical axis represents test loss. A model positioned further to the upper right indicates better performance. (**B**) ROC curve of RCN showing the true positive and false negative rates, with the AUC value calculated for testing images. (**C**) Confusion matrix of the RCN classification results.

All models achieved classification accuracy more than 75%. The top three models, based on both accuracy and test loss, were RCN, Xception, and EfficientNet, all of which exhibited classification accuracies above 82% and test losses below 0.076. Notably, RCN had the lowest test loss, at less than 5%, suggesting superior generalization performance for mouse images. Additionally, the ROC-AUC of RCN was 0.9 (**Fig. 4B**), demonstrating its practical classification capability. Furthermore, the confusion matrix (**Fig. 4C**) emphasizes that even with the quality limitations the model classifies efficiently.

To examine which regions of mouse images were utilized by the proposed RCN model for classification, we computed saliency map. Three mice were randomly selected from both the correctly classified estrous (**Fig. 5A**) and non-estrous groups (**Fig. 5B**) for comparison. The saliency map revealed that highlighted areas extended from the vaginal region toward the anus, while the vaginal opening itself was not specifically emphasized. Interestingly, the vaginal opening itself was not particularly highlighted; instead, the model utilized features from the entire vaginal area extending to the anus, rather than relying solely on the vaginal region for classification which indicates that the model uses not only the vaginal area as a parameter.

**Figure 5.**
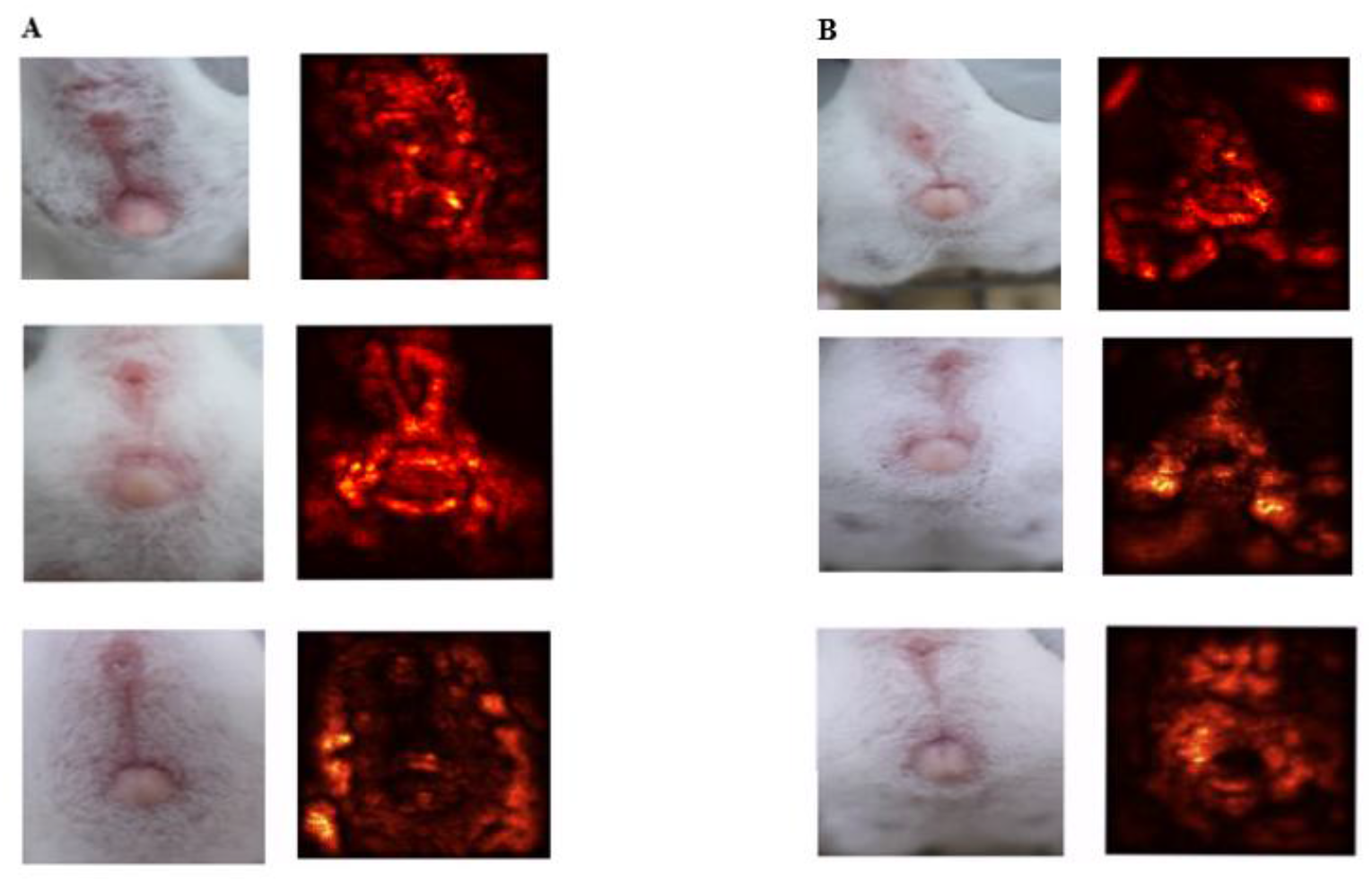
Visualization of pixel importance using saliency map generated by the RCN model. (**A**) Images from the estrus class. (**B**) Images from the non-estrus class.

## Discussions

This study presents a deep learning-based approach for binary classification of the estrous cycle in female mice using external genital images. Our proposed model, Repro Cycle Net (RCN), achieved a classification accuracy of over 83% and demonstrated the lowest test loss among all tested models, despite its relatively simple four-layer architecture. The low-test loss indicates high confidence in prediction outcomes and suggests robust generalization with almost no overfitting. These results highlight the novelty and practicality of RCN, which achieves competitive performance while maintaining a lightweight and simple architecture.

The lightweight design of RCN offers distinct advantages in computational efficiency, enabling implementation in resource-limited settings such as long-term monitoring systems or on-site inference in laboratory animal facilities. Furthermore, unlike large-scale pretrained models, RCN allows greater flexibility in optimization and reduced risk of overfitting, making it suitable for specialized tasks.

In comparison to previous approaches, our model also demonstrates a distinct contribution in both methodology and practical utility. Previous studies on estrous cycle classification have primarily relied on cytological images derived from vaginal smears, which are often invasive, labor-intensive, and require technical expertise. While several machine learning models have been developed for this task, most have utilized complex architectures or required high-resolution microscopic inputs. In contrast, our study introduces a classification model based solely on macroscopic, non-invasive external images, offering a more accessible and animal welfare– conscious alternative. The proposed RCN achieves a competitive accuracy without relying on pretrained networks or extensive model depth, indicating that effective classification does not necessarily require complex model structures when appropriate feature representations are learned. To our knowledge, few if any prior studies have reported comparable performance using external genital images alone, underscoring the originality of this approach.

From an experimental biology perspective, the ability to accurately and consistently identify estrous stages from external images has direct implications for improving experimental design and reproducibility. Hormonal fluctuations during the estrous cycle can significantly influence physiological, behavioral, and developmental outcomes. Thus, misclassification or neglect of cycle stages can introduce confounding factors into experimental results. By providing a rapid and objective tool for cycle staging, our model supports more rigorous data interpretation, especially in longitudinal or high-throughput experimental setups. Furthermore, owing to its low computational demand, the model can be deployed locally on standard laboratory equipment without requiring GPU resources, making it feasible for routine use in animal facilities. These technical and biological advantages highlight the broad potential utility of the model in various research environments. Importantly, a binary classification of estrous versus non-estrous is sufficient to guide mating decisions, directly supporting reproductive studies and colony management. This practical applicability highlights that even without four-stage classification, the model provides immediate benefits to experimental workflows.

Saliency map analyses revealed that the model predominantly focused on the vaginal and anal regions, while the vaginal opening itself was not strongly emphasized. White fur regions were largely disregarded, and the model appeared to rely on features derived from the exposed perivaginal area. Although no quantitative color analysis was performed, it is possible that the model utilized subtle differences in brightness or hue—potentially in the reddish range—between the prevaginal and perianal areas. This suggests that the model may have captured discriminative visual cues not typically recognized in conventional human assessments. Nevertheless, in rare cases the model still produced misclassifications, as observed in the confusion matrix (**Fig. 4C)** and certain saliency map examples (**Fig.5A, 5B**), indicating the need for further optimization and refinement.

To ensure label consistency and minimize noise, visual assessments were performed with the assistance of domain experts. Given the subjectivity inherent in visual staging, this expert involvement was crucial for establishing reliable ground truth labels, which likely contributed substantially to improving model performance and training stability.

## Limitations and Future Directions

Several limitations should be acknowledged. First, the quality and consistency of image acquisition have a critical impact on classification performance. Key features such as vaginal coloration or opening may be obscured due to inconsistent lighting or fur coverage. Future studies may benefit from standardized illumination and careful consideration of ethically feasible fur removal or trimming to improve visibility of relevant features.

The present dataset was limited to albino ICR mice. Extending the dataset to include additional strains with diverse genetic backgrounds and pigmentation will be necessary to improve model generalizability. Moreover, the current model addresses only binary classification, whereas the biological estrous cycle comprises four stages. Expanding the classification to all four stages will require a substantially larger and more finely annotated dataset, although binary classification may remain practical for many experimental applications where the presence or absence of estrogenic activity is the primary concern.

Further improvements may also be achieved through standardized animal handling procedures during imaging and stratification by age to account for physiological variability. Additionally, future work will explore ensemble modeling strategies and incorporate a broader set of performance metrics to further enhance robustness and accuracy.

## Authors contribution

- Conceptualization and study design: YS
- Data collection and preprocessing: YT, MS
- Model implementation and analysis: YT, MS
- Supervision of experiments / Ethical compliance: AI, GPS
- Manuscript writing and revision: YT, MS, YS
- Final approval of the manuscript: YS, AI

## Acknowledgements

We would like to thank Kenta T. Suzuki for his assistance with hardware preparation and helpful advice on programming. This research was supported by the Organ Developmental Engineering Laboratory within the Division of Biological Sciences at the Nara Institute of Science and Technology, Japan, during the period of 2022–2023. Additional support was provided by the Data Science Center at the Nara Institute of Science and Technology through the Grants for New Issue Identification Activities program.

